# Aberrant perceptual judgements on speech-relevant acoustic features in hallucination-prone individuals

**DOI:** 10.1101/2020.06.26.171330

**Authors:** Julia Erb, Jens Kreitewolf, Ana P. Pinheiro, Jonas Obleser

**Affiliations:** Department of Psychology, University of Lübeck, Lübeck, Germany; Center for Brain, Behavior, and Metabolism, University of Lübeck, Lübeck, Germany; Faculdade de Psicologia, Universidade de Lisboa (FPUL), Lisbon, Portugal

**Keywords:** Speech perception, psychoacoustics, reverse correlation, spectro-temporal modulations, auditory verbal hallucinations, schizotypy

## Abstract

Hallucinations constitute an intriguing model of how percepts are generated and how perception can fail. Here, we investigate the hypothesis that an altered perceptual weighting of the spectro-temporal modulations that characterize speech contributes to the emergence of auditory verbal hallucinations. Healthy adults (N=168) varying in their predisposition for hallucinations had to choose the ‘more speech-like’ of two presented ambiguous sound textures and give a confidence judgement. Using psychophysical reverse correlation, we quantified the contribution of different acoustic features to a listener’s perceptual decisions. Higher hallucination proneness covaried with perceptual down-weighting of speech-typical, low-frequency acoustic energy while prioritising high frequencies. Remarkably, higher confidence judgements in single trials depended not only on acoustic evidence but also on an individual’s hallucination proneness and schizotypy score. In line with an account of altered perceptual priors and differential weighting of sensory evidence, these results show that hallucination-prone individuals exhibit qualitative and quantitative changes in their perception of the modulations typical for speech.

**Author summary:** Hallucinations -- that is, percepts in the absence of an external stimulus -- are
prevalent in psychotic disorders such as schizophrenia, but also occur in the general
population. To date it is unknown whether the emergence of hallucinations is rooted in an altered perception of sounds. Fusing the psychophysical technique of reverse correlation with concepts from computational psychiatry, this research reveals alterations of sensory processing in hallucination-prone adults. We show that the higher nonclinical adults’ predisposition to hallucinations, the more they prioritise the sound features atypical for speech such as higher frequencies. At the same time, they express higher confidence in their perceptual judgements. The present approach may contribute to improving early diagnosis and prevention strategies in individuals at risk for psychosis.

## Introduction

A major challenge of sensory neuroscience remains to understand how adaptive top-down weighting of sensory evidence due to, e.g., ongoing task demands influence percepts. As hallucinations occur in the absence of an external stimulus, they constitute an intriguing model for the generation of percepts. Hallucinatory experiences, mostly visual or auditory, are prevalent in psychotic disorders such as schizophrenia, but also have an estimated prevalence of 6-13% in the general population (‘non-clinical voice hearers’) ^1,2^, consistent with the hypothesis that psychosis exists on a continuum with normal experience ^3^. Auditory (but not visual) perceptual abnormalities predict conversion to psychosis in individuals at risk of psychosis^4^.

Psychosis is thought to result in part from aberrant integration of prior knowledge with incoming sensory information. Accumulating evidence suggests strong prior expectations to play a critical role in the emergence of hallucinations ^5^. This evidence may first appear at odds with predominant accounts of weaker priors in hallucination proneness and psychosis. However, hierarchical predictive processing frameworks can account for evidence of both weaker and stronger priors, but at different levels of a hierarchy of information processing^5,6^.

At some levels of the information processing hierarchy, individuals prone to psychosis appear to rely less on prior knowledge than on the sensory information. One prominent example is the resistance to the hollow mask illusion in people with schizophrenia spectrum disorders^7^.

Several observations, however, imply an increased bias towards top-down information in hallucination proneness. For example, prior knowledge of an image confers an advantage in recognizing that image when it is degraded; individuals at risk of psychosis are more susceptible to this perceptual advantage ^8^. Notably, hallucination-prone participants performed better in the recognition of both the image gist and the details by relying on prior knowledge ^9^. Further, a recent visual-auditory conditioning study^10^ used a visual conditioned cue to predict a faint auditory stimulus. While all participants experienced conditioned auditory hallucinations when presented with solely the visual stimulus, hallucination-prone individuals were more susceptible to such conditioned hallucinations ^10^. For speech, a similar effect has been reported: Voice-hearers listening to degraded (sine-wave) speech, showed stronger expectations to hear speech than controls who recognized the presence of speech later when not explicitly instructed ^11^. At higher processing levels, stronger semantic expectations have been reported as well in hallucination proneness. In a speech-in-noise recognition task where predictability of the final word in a sentence was manipulated, hallucination-prone individuals showed a stronger tendency towards hearing a predictable word that fits the context^12,13^. These results are commensurate with a bias towards top-down processing at various levels of the processing hierarchy which may contribute to hallucinatory experiences^14^.

Regarding the neurobiology of auditory verbal hallucinations (AVH), functional MRI studies report activation of auditory cortex ^15^, including primary auditory cortex ^15^, when patients experience AVH (for review see ^17^). To date, it is unclear in how far the general response properties of auditory cortex are altered in voice hearers. A recent strain of research provides converging evidence that the healthy human auditory cortex analyzes sounds along so-called spectro-temporal modulations: The auditory pathway is thought to not only implement forms of “tonotopic” frequency analysis^18^, but to rather represent sound as frequency-specific spectral and temporal modulation filters ^19,20^ for neurobiological evidence see e.g., ^21,22^). Hallucinations in schizophrenia have been linked to deficits in object formation ^23^. Auditory object formation in turn is known to rely heavily on the extraction of the spectro-temporal modulations in the auditory scene^24^.

In the present study we therefore examine whether differences in the perception of these spectro-temporal modulations abundant in speech ^25^ can be linked to a propensity towards auditory hallucinations. We here investigate processing of spectro-temporal modulations in individuals presenting with varying, non-clinicai degrees of predisposition to hallucinations.

First, we establish individual listeners’ “*speechiness* kernels”, that is, an individual template of those acoustic features that elicit a speech percept. To this end, we present two ambiguous sound textures in noise^26^ to human listeners and ask them to the ‘more speech-like’ one. This allows us to retrieve their internal representation that drives the categorization into speech, using the psychophysical technique of reverse correlation ^27,28^. Second, we relate those speechiness kernels to the degree of individual schizotypal traits (subscale ‘unusual perceptual experiences’ ^29,30^), and to individual hallucination proneness ^31,32^, probing a psychosis continuum. Compared to studies with psychotic patients, the study of non-clinicai participants has the important advantage to circumvent confounding factors such as medication and presence of other symptoms (e.g., negative symptoms^33^).

The results pose an intriguing link between current models in computational psychiatry and recent advances in modelling the perceptual and neural response in auditory neuroscience.

## Results

In a short online (*N* = 131) and an extended lab version (*N* = 37) of a 2-AFC experiment with confidence judgement, participants had to choose the ‘more speech-like’ of two presented ambiguous sound textures (Fig. 1). Using reverse correlation, we obtained perceptive fields termed *“speechiness* kernels” that quantify the contribution of different acoustic features to a listener’s perceptual decisions. Hallucination proneness was assessed with the Launay-Slade hallucination scale (LSHS). Additionally, we evaluated schizotypy using the Schizotypal Personality Questionnaire (SPQ) with a particular interest in the subscale “unusual perceptual experiences” (SPQ-UP) as second measure of predisposition to unusual perception.

**Fig. 1.**
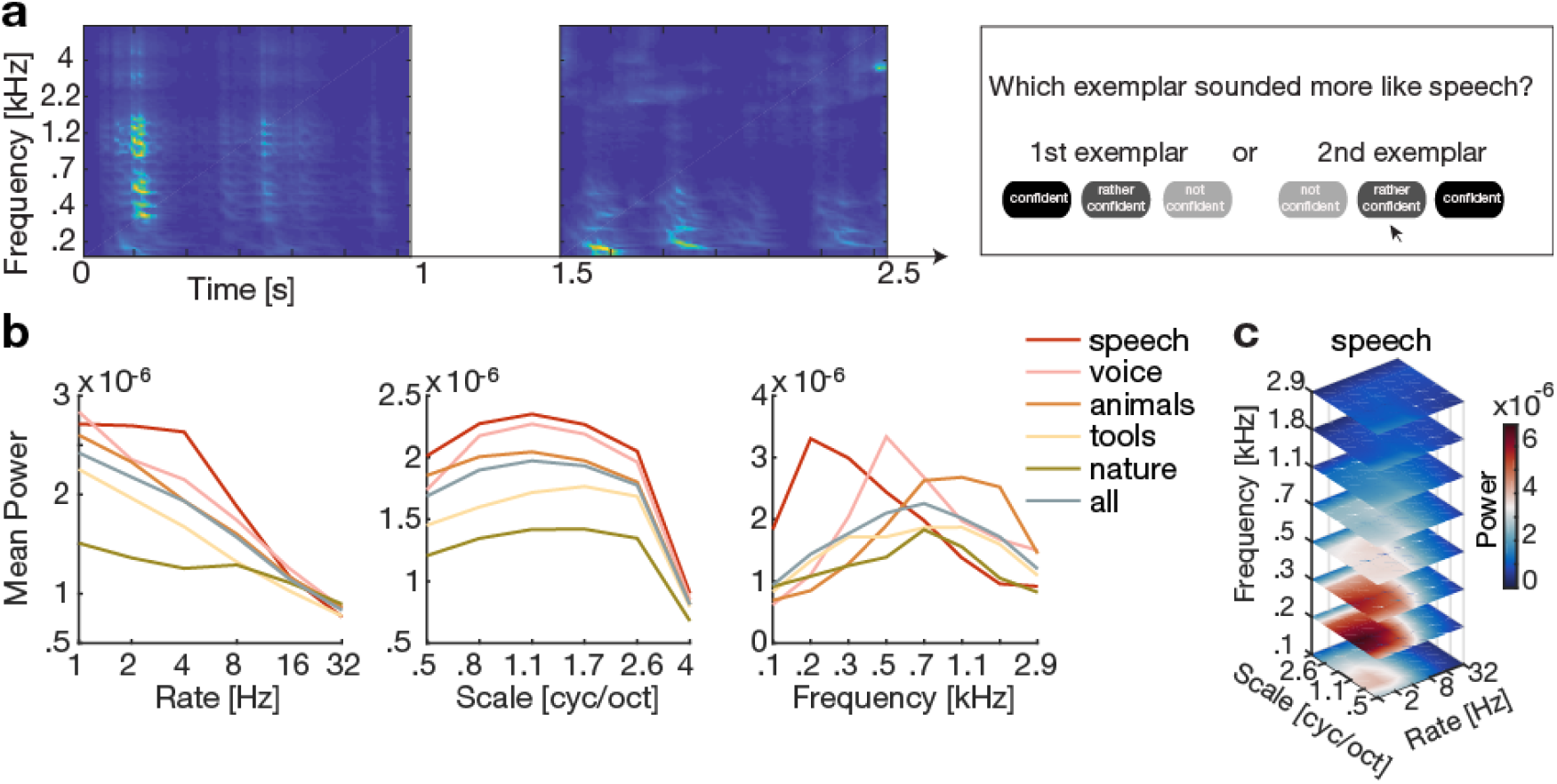
Stimuli and task. (a) In a 2-alternative-forced-choice (2-AFC) task with confidence judgement, participants were presented with two sound textures (here shown as spectrograms) and were asked to simultaneously express their decision and confidence about which exemplar sounded more like speech, (b) Modulation spectra for sound textures, marginalized for each acoustic dimension (rate, scale, frequency) and split by categories of the original sounds from which the textures were synthesized (see legend), (c) Average modulation spectrum for the speech textures.

### Schizotypal traits and hallucination predisposition

In the lab experiment, LSHS scores ranged from 1 to 32 (median = 10: max possible score 48) and global SPQ scores ranged from 3 to 37 (median = 16, max possible score 74). In the online experiment, LSHS scores ranged from 1 to 33 (median = 9) and global SPQ scores ranged from 0 to 41 (median = 15: see Fig. 2a).

**Fig. 2.**
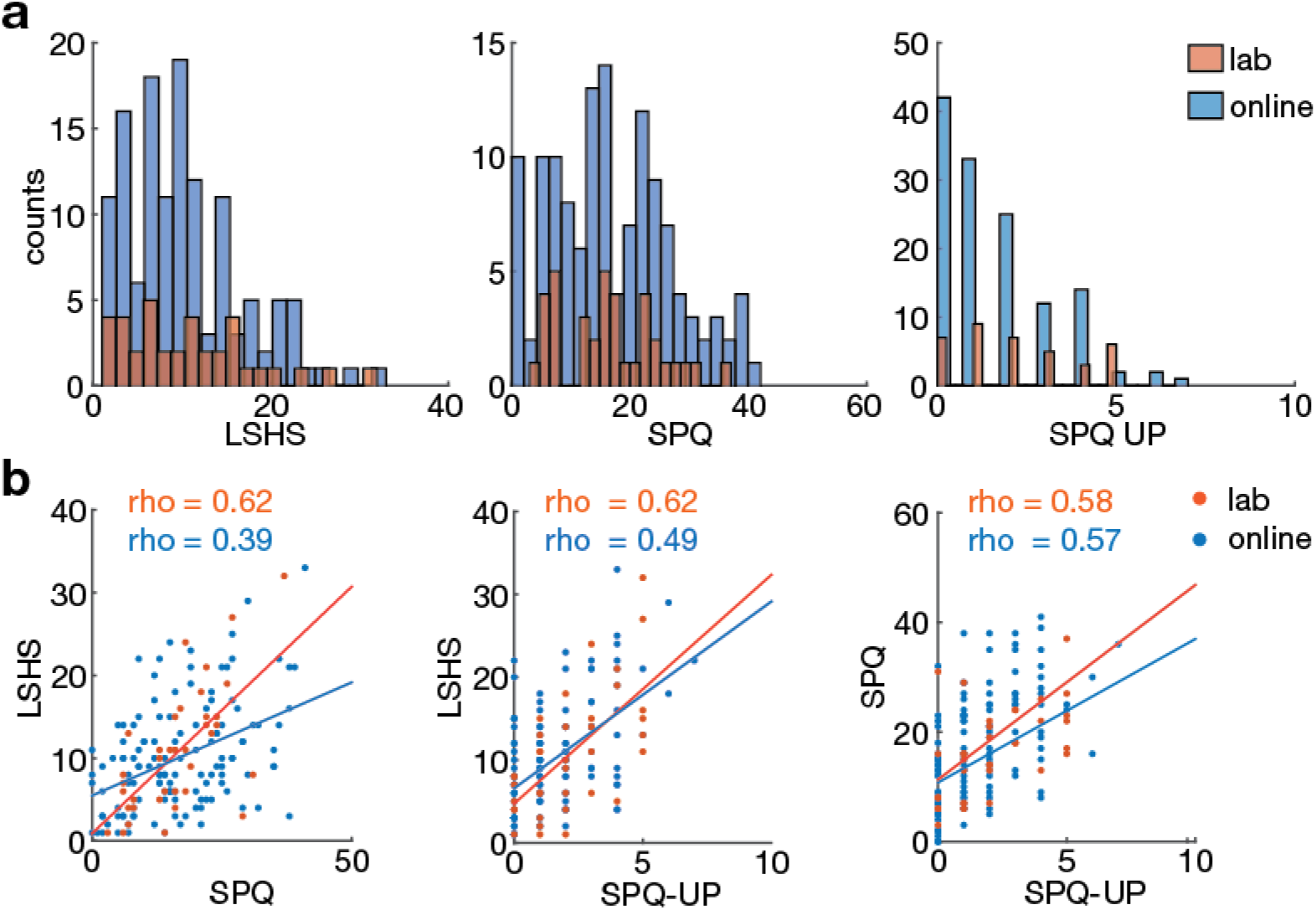
Schizotypy and hallucination scale results. (a) Histograms for LSHS and SPQ scores separately for lab (orange) and online (blue) experiment. Note that the maximum possible score is 42 for LSHS, 74 for SPQ, and 9 for the subscale SPQ-UP. (b) Scatter plots showing the correlations between LSHS, SPQ total score and SPQ-UP. Spearman’s correlation coefficients are shown separately for lab (orange) and online experiment (blue). All scales are significantly correlated (*p* < 0.001), *R*^2^ ranges from .18 to .42. LSHS: Launay-Slade hallucination scale; SPQ: schizotypal personality questionnaire; SPQ-UP: subscale unusual perceptual experiences.

Across both experiments (*N* = 168), LSHS scores were uncorrelated with age (*r* = −.098, *p* = .206) or gender (*r* = −.077, *p* = .321). The same held for global SPQ scores (age *r* = .099, *p* = .201; gender *r* = .140, *p* = .070). More importantly, LSHS scores, global SPQ scores and the subscale SPQ-UP were all substantially correlated amongst each other (Fig. 2b). The intercorrelations of 18-42 % shared variance emphasise the convergent validity of the questionnaires used to measure individual predisposition to unusual perceptual experiences here.

To examine potentially differential effects of hallucinations and delusions, we looked at the SPQ subscale ‘Magical Thinking’ (SPQ-MT; supplementary Fig. S1a). SPQ-MT scores were significantly correlated with LSHS scores (supplementary Fig. S1b), underlining the common aspects between both scales measuring predisposition to delusions and hallucinations, respectively.

### Speechiness kernels

First, we analyzed how participants’ judgements of speechiness varied as a function of the spectro-temporal modulations contained in the sound textures. To obtain such speechiness kernels, we used reverse correlation by contrasting the averaged spectro-temporal modulations of the stimuli judged as more versus less speech-like. Speechiness kernels averaged across participants were highly correlated between the lab and online studies (Fig. 3a).

**Fig. 3.**
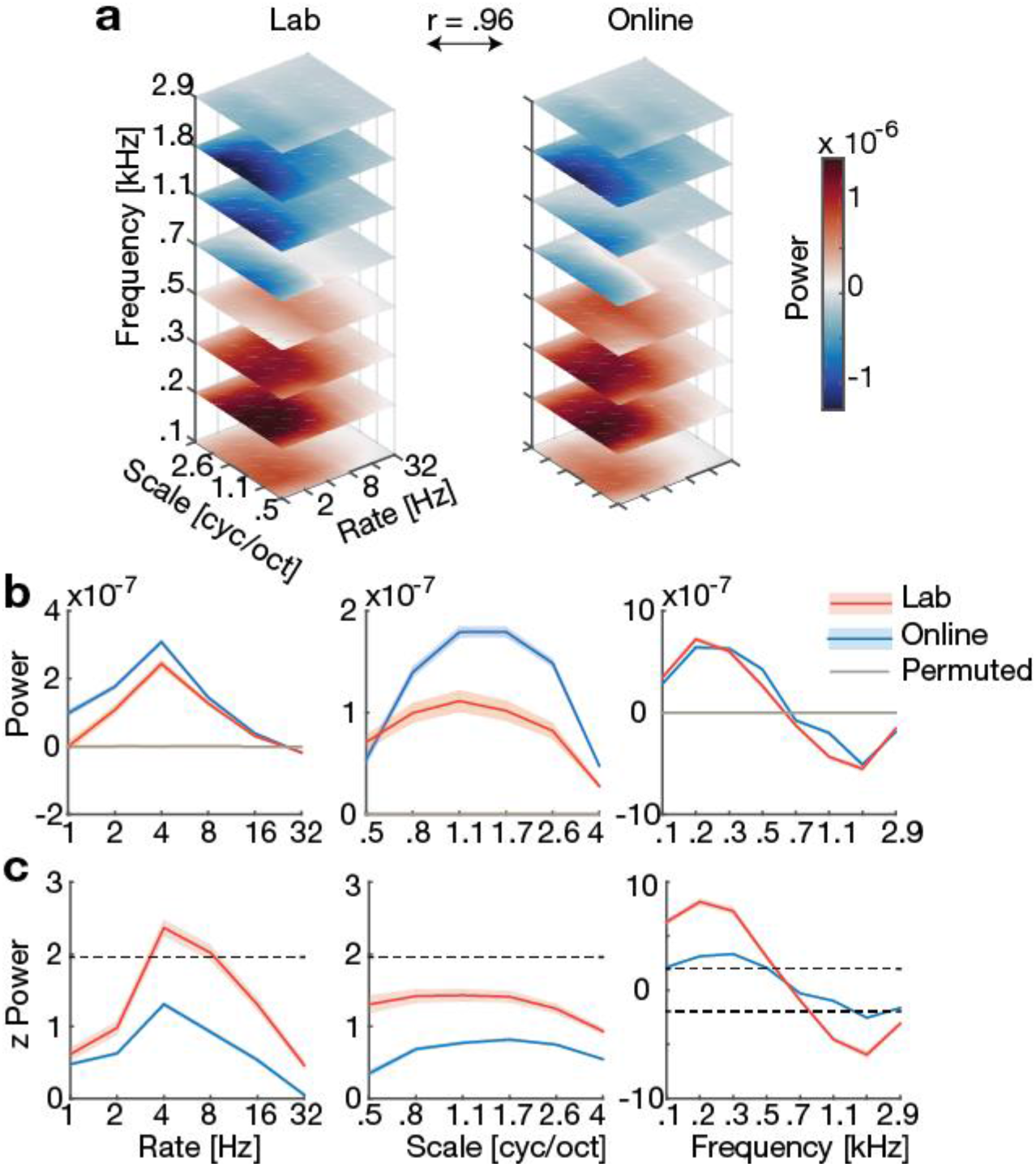
Speechiness kernels. (a) Averaged speechiness kernels for lab and online study are highly correlated (Pearson’s *r).* (b,c) Marginal profiles of speechiness kernels (mean ± standard error) for lab and online study separately before (b) and after (c) z-scoring relative to the empirical null distribution (obtained with N = 10,000 permutations). Absolute z-scores exceeding 1.96 are considered significant (indicated by the dashed line). Due to the markedly lower number of trials in the online (*n* = 108) compared to the lab study (*n* = 540), the unstandardized power of the speechiness kernels is higher in the online study (b). However, after z-scoring the speechiness kernels, the lab exceeds the online kernel, reflecting higher precision in the lab measurements (c).

The marginal profiles expectedly peaked for frequency at 200 Hz and for temporal modulations at ~4 Hz, indicating high speechiness judgements when acoustic power was high at low frequencies and slow temporal modulations (Fig. 3b). As an outlook, those peaks were driven by the trials on which the participants were confident (see Fig. 5a). We permuted participants’ responses (n = 10,000 permutations) to obtain the empirical null distribution of the speechiness kernels (Fig. 3b, yellow line) and *z*-scored empirical kernels relative to the null distribution. *Z*-scores proved significant (i.e., |*z*| > 1.96) for temporal rates of 4–8 Hz in the lab experiment, and for low and high frequencies in both the lab and online experiment (Fig. 3c). These findings are in line with numerous studies showing the importance of those features to speech perception ^25,34-36^, confirming the feasibility of the current method to obtain meaningful perceptive fields for speech.

#### Kernel stability

In the lab experiment, average speechiness kernels based on the first 100 trials were highly similar to the total kernel (i.e., comprising all 540 trials, mean [SE] Pearson’s *r* = 0.926 [0.026], see supplementary Fig. S2). We took this finding as evidence that the lower number of 108 trials used in the online experiment suffice to obtain a stable estimate of the speechiness kernel.

### Relation of acoustic feature weighting to hallucinatory predispositions

Next, we asked whether the individual extent of hallucination proneness is related to particular features of the speechiness kernel. First, we conducted a principal component analysis (PCA) to reduce the dimensionality of the speechiness kernels from 288 components (one for each feature of the speechiness kernel) to the first six components (see Methods). This procedure was justified by the drop in eigenvalues after the sixth component (see scree plot Fig. 4a). There was one clear dominant component (component 1) and two minor components (component 2 and 3; whose eigenvalue still clearly exceeded the eigenvalues of surrogate data, Fig. 4a). The PCA approach also held the advantage of yielding, by design, independent regressors to be subsequently used in a linear model. The first component was characterized by overall uniformly distributed high energy except in some high frequency bins (Fig. 4b blue line, Fig. 4d, left). The third component resembled a typical speechiness kernel (see Fig. 4c for the average kernel), exhibiting higher energy at the lower frequencies (Fig. 4b, Fig. 4d).

**Fig. 4.**
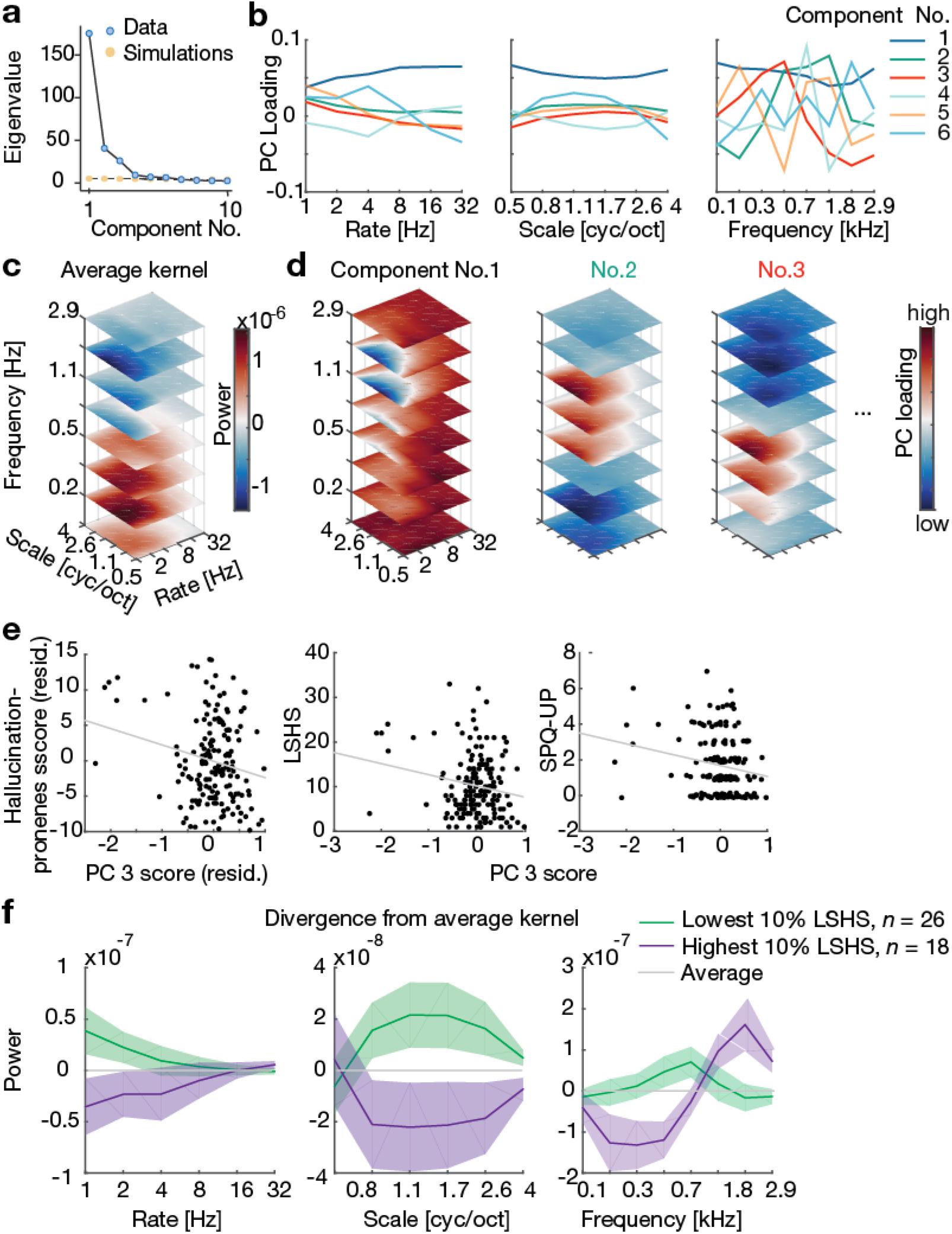
Principal component analysis (PCA) of the speechiness kernels. (a) Scree plot of the PCA performed jointly on all data (online and lab experiment). For subsequent analyses, only the first six components were retained because for all further components the eigenvalue (blue) was smaller than eigenvalues based on randomly generated data (yellow). There was one clear dominant component (component 1) and two minor components (component 2 and 3). (b) Marginal profiles of the loadings on the first six components. Note the low-frequency energy peak in component 3 (red), (c) Speechiness kernel averaged over all participants and experiments (see also Fig. 3). (d) Loadings for the first three components in the modulation space, (e) Hallucination proneness scores (the average of *z*-scored LSHS and SPQ-UP scores) were regressed against the first six components. The residuals of this compound hallucination proneness score and the third component from this multiple regression (Table 1) are shown in the scatter plot (left). Scores of the third component as a function of LSHS (middle) and SPQ-UP (right) are shown in scatter plots. LSHS and SPQ-UP scores are correlated to scores of the third component only. For display purposes only, the integer SPQ-UP scores were jittered slightly by adding a uniformly distributed random quantity between −0.15 and +0.15. (f) To illustrate the LSHS effects, the differences in the marginal profiles of the speechiness kernel relative to the average kernel are shown for the participants within the lowest (*n* = 26, green) and highest decile (*n* = 18, violet) of LSHS scores. Note that the most hallucination-prone participants (violet) tend to overweight high frequencies while downweighting low frequencies (left plot). Error bands show standard error of the mean (SE). PC: Principal Component; LSHS: Launay-Slade hallucination scale; SPQ-UP: schizotypal personality questionnaire subscale unusual perceptual experiences.

To capture the common aspects of both questionnaires, we averaged individual *z*-scored LSHS and SPQ-UP scores into a compound hallucination proneness score. We used individual component scores of the first six components to predict hallucination proneness scores in a multiple regression analysis (Table 1). To account for outliers (cf. Fig. 4e) we performed a robust multilinear regression ^37^. The hallucination proneness score was significantly related to the scores of the third component (Table 1, Fig. 4e), surviving false discovery rate (FDR) correction for multiple comparisons.

**Table 1:** Robust Multiple Linear Regression predicting a compound hallucination proneness score (average of z-scored LSHS and SPQ-UP scores) based on the first six principal component (PC) scores of the individual speechiness kernels and a regressor of no interest coding experiment type based on data from *N* = 168 individuals. Shown are *β*-estimates, standard error of coefficient estimates (SE), t-values and p-values for the robust regression. Note that for the least squares solution of the same equation *R*^2^ = 0.095.

| Hallucination proneness score |  |  |  |  |
| --- | --- | --- | --- | --- |
| Predictors | $\beta$ | SE | t-value | p-value |
| PC No.1 | -0.0263 | 0.0607 | -0.4333 | 0.6654 |
| PC No.2 | -0.0479 | 0.0985 | -0.4866 | 0.6272 |
| **PC No.3** | **-0.3523** | **0.1282** | **-2.7489** | **0.0067** |
| PC No.4 | -0.4136 | 0.2118 | -1.9528 | 0.0526 |
| PC No.5 | -0.2274 | 0.2196 | -1.0359 | 0.3018 |
| PC No.6 | 0.3398 | 0.2716 | 1.2512 | 0.2127 |
| Experiment<br>(lab \| online) | -0.1207 | 0.0825 | -1.4638 | 0.1452 |

LSHS and SPQ-UP scores separately also predicted how a listener would perceptually weight scores of the third principal component (PC3), which was dominated by low-frequency acoustic energy (Fig 4b, d). PC3 scores covaried negatively with an individual’s tendency towards aberrant perception, that is, both the LSHS (Pearson’s *r* = −0.192, *p* = 0.013, Spearman’s *rho* = −0.066, p = 0.393, Mutual Information *MI* = 0.010, p = 0.063, Bayes factor BF_10_ = 2.126) and SPQ-UP score (*r* = −0.195, *p* = 0.011, rho = −0.144, *p* = 0.062, *MI* = 0.028, *p* = 0.006, BF_10_ = 2.306; Fig. 4e; see also Table 1).

These results provide evidence for an association of higher hallucination proneness with reduced reliance on the speech-typical low frequencies for classifying stimuli into speech (Fig. 4b, d, f). This effect is illustrated in the markedly different speechiness kernels of the individuals with the lowest and highest 10% LSHS scores: Hallucination-prone participants tend to prioritise high frequencies and downweight low frequencies relative to the average kernel when categorizing stimuli into speech (Fig. 4f, violet).

The subscale SPQ-MT also covaried with PC3 scores (Supplementary Fig. S1c), indicating that propensity to delusions was similarly associated with atypical internal templates for acoustic features in speech.

### Sound discriminability

Human fMRI studies suggest that spectro-temporal population tuning of auditory cortex maximizes the acoustic distance between speech sounds, facilitating our ability to discriminate speech^22^. Under the assumption that the speechiness kernels reflect perceptive fields of speech, we sought to investigate (1) how individual speechiness kernels affect discriminability of presented stimuli and (2) whether this differs in hallucination proneness.

We expected that the speech-like modulation content in a sound texture should be a prominent driver of the listeners’ speechiness judgements. Therefore, we looked at discriminability of sound pairs presented on each trial (i.e., the sensory evidence). To test how individual speechiness kernels shape this discriminability, we weighted the sounds by individual kernels.

First, we projected sounds through the individuals speechiness kernels in the modulation representation. We calculated the Euclidian distance between original sound pairs and projected sound pairs presented on each trial (see Materials and Methods). A comparison between original and projected sound pair distance showed increased discriminability of sound pairs after projection through individual kernels (Fig. S3a).

Second, to quantify benefits from individual speechiness kernels, we fitted individual linear slopes to projected as a function of original sound-pair distance. Individual slopes (“speechiness kernel benefit”) were significantly higher than one (*t*(167) = 32.61, *p* < .001), indicating the “warping” of an acoustic into a perceptual distance representation and validating the present speechiness kernel approach. Notably, however, individual speechiness kernel benefits were unrelated to LSHS scores *(r* = −0.045, *p* = 0.559, Fig. S3b) with evidence for the absence of an effect as indicated by the Bayes Factor (BF_01_ = 8.749). These results suggest that internal templates for speech amplify the discriminabiiity of sound textures, irrespective of an individual listener’s hallucination proneness.

### Confidence judgements

To investigate whether confidence affects the speechiness kernels, we calculated three different kernels, one for each confidence level separately (Fig. 5a). Although the shape of the kernels was similar for all confidence levels, higher confidence amplified the magnitude of the kernel (see also Fig. 5b). The proportion of “confident” responses was positively correlated both with the LSHS scores and global SPQ scores, indicating that participants with higher hallucination proneness and schizotypy reported more often that they were confident about their speechiness judgements (Fig. 5d), This result is consistent with previous evidence for hallucination-prone individuals expressing more confidence in their decisions ^10,38^ (for review see ^39^).

**Fig. 5.**
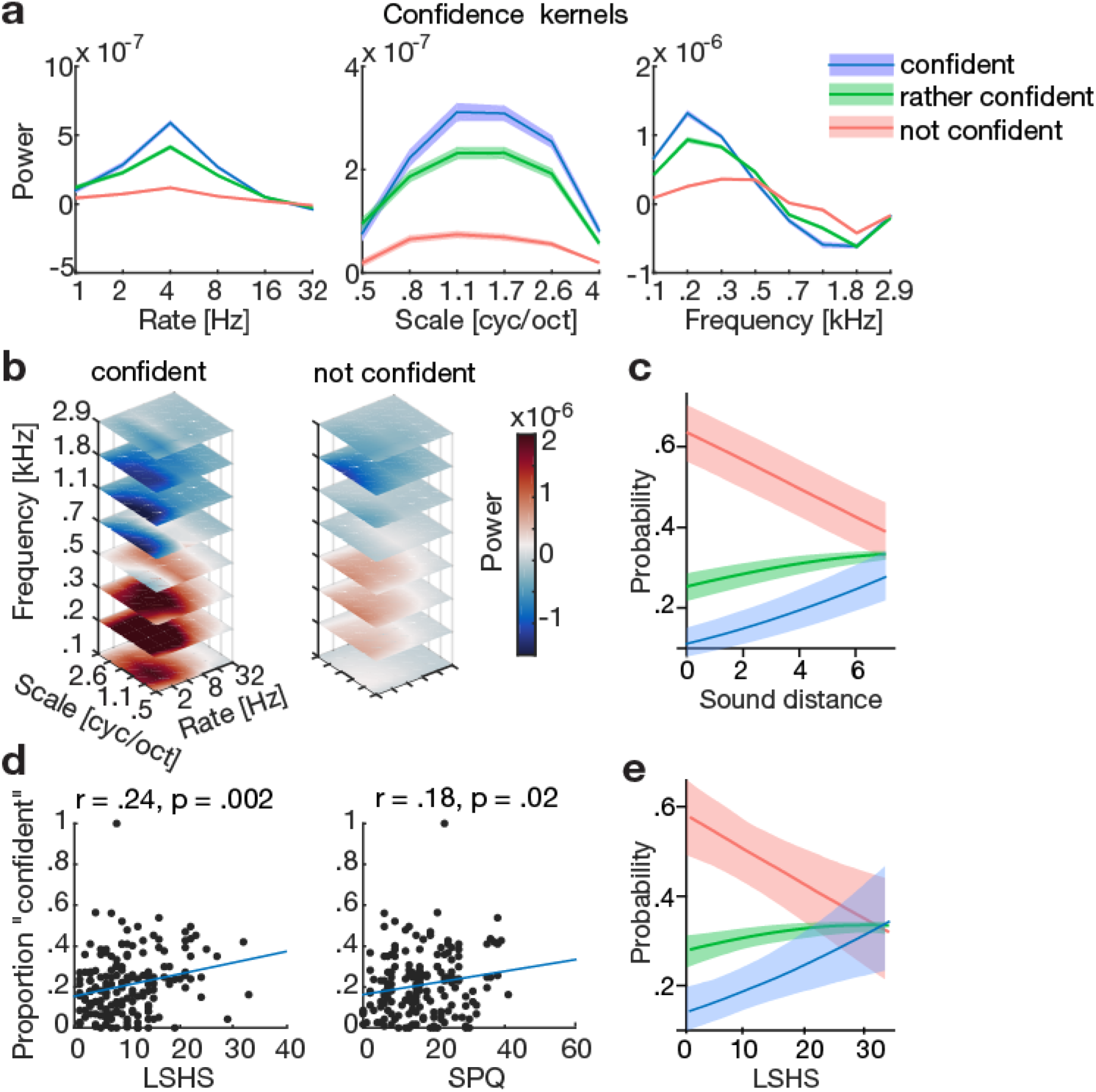
Confidence judgements in speechiness kernels (jointly from lab and online experiment) and relation of hallucination proneness to confidence judgements. (a) Marginal profiles of speechiness kernels (mean ± SE) for three different confidence levels separately, (b) Speechiness kernels for the two extreme confidence levels (confident, not confident), (c) Estimates from a generalized linear mixed model (GLMM) predicting confidence levels based on sound pair distance; the model also has the predictors LSHS, SPQ and experiment type (see below); same color legend same as in (a), (d) Proportion of “confident” judgements correlated (Pearson’s r) with LSHS and global SPQ scores, (e) Estimates from a GLMM predicting confidence levels based on LSHS, model also has the predictors sound distance, SPQ and experiment type (see below); same color legend same as in (a). LSHS: Launay-Slade hallucination scale, SPQ: schizotypal personality questionnaire.

A more mechanistic explanation of this correlation would afford that this relation of confidence judgements to hallucination proneness also holds at the trial-by-trial level, where we can account for stimulus discriminability, experiment type, and subject-specific intercepts.

Using ordinal linear mixed-effects regression, we regressed trial-by-trial confidence judgements (on a Likert scale from 1 [“not confident”] to 3 [“confident”]) against the following predictors: trial-wise projected sound pair distance (pairwise Euclidian distance of the sounds in the modulation representation filtered by individual speechiness kernels); the hallucination proneness variables LSHS score; experiment (a binary indicator variable coding online versus lab); the interaction sound-pair distance x experiment; and a subject-specific random intercept. In a second model we used the same predictors but replaced the LSHS scores with the global SPQ score. Note that we used projected rather than original sound pair distance to see how confidence emerges from the participants’ perceptions of the stimuli (cf. discriminability analysis above). The null model excluded hallucination proneness variables thus comprising the following predictors: projected sound pair distance; experiment (coding lab versus online); the interaction projected sound-pair distance x experiment; and a subject-specific random intercept.

Greater projected sound-pair distance, but also higher LSHS and global SPQ scores were significantly associated with higher confidence judgements (Table 2; Fig. 5c, e). The interaction of sound pair distance with experiment type was a significant predictor of confidence judgement when considering a 95% credible interval, indicating that the confidence judgement depended more on sensory evidence in the lab than in the online study. Observing these data was about thirteen times more likely under a model including the LSHS score than under a null model with the same parameters except LSHS, as evidenced by an average Bayes Factor BF_LSHS–null_ of 13.07, 95% Cl [12.00; 14.13], Echoing the convergent validity of LSHS and SPQ, global SPQ score also proved a significant predictor of confidence judgement: a very comparable magnitude was observed for the Bayes Factor of a model including the global SPQ score relative to a null model with the same predictors except SPQ (BF_SPQ–null_ of 12.68, 95% Cl [11.93; 13.43]).

**Table 2:** Generalized Linear Models. (Ordinal regression) predicting confidence judgements on a Likert scale from 1 (“not confident”) to 3 (“confident”). Shown are estimates of effects of interest as Log Odds with a 95% Bayesian highest posterior density interval (labelled “95% Cl”, credible interval). The models entailed data from a total of 34,128 single trials from N=168 participants. The two models only differ in the predictor LSHS or global SPQ score, respectively. The Null model includes the same predictors except LSHS or global SPQ scores. Note that predictors are considered significant when the 95% credible interval dœs not include zero (marked in bold).

| Confidence Judgement |  |  |  |  |  |
| --- | --- | --- | --- | --- | --- |
| *Predictors* | *Log-Odds* | *CI (95%)* | *Predictors* | *Log-Odds* | *CI (95%)* |
| Intercept 1 | 0.00 | -0.16 – 0.16 | Intercept 1 | 0.00 | -0.18 – 0.16 |
| **Intercept 2** | **0.88** | **0.71 – 1.04** | **Intercept 2** | **0.87** | **0.70 – 1.04** |
| **LSHS** | **0.15** | **0.07 – 0.23** | **Global SPQ** | **0.13** | **0.06 – 0.20** |
| **Projected sound distance** | **0.11** | **0.10 – 0.13** | **Projected sound distance** | **0.11** | **0.10 – 0.13** |
| Experiment (lab \| online) | -0.10 | -0.28 – 0.09 | Experiment (lab \| online) | -0.11 | -0.31 – 0.09 |
| **Projected sound distance x Experiment** | **-0.10** | **-0.13 – -0.08** | **Projected sound distance x Experiment** | **-0.10** | **-0.13 – -0.08** |
| **Random Effects** |  |  |  |  |  |
| $\sigma^2$ | 0.03 | | | | |
| $\tau_{00}$ | 0.59 | | | | |
| ICC | 0.04 |  |  |  |  |

## Discussion

In how far does hallucination proneness in non-clinical participants manifest in the aberrant perceptual judgement of acoustic features, namely spectro-temporal modulations? We studied this using a simple “speechiness” judgement with confidence ratings based on synthesized sound textures, both in a short online and an extended lab experiment and gathered a total of N=168 data sets.

First, we found higher scores on both the schizotypy personality questionnaire (SPQ) subscale ‘unusual perceptual experiences’ and on the Launay-Slade Hallucinations Scale (LSHS) to covary with the degree to which they classified textures as ‘speech’ that were lacking the speech-typical low-frequency dominance.

Second, those individuals scoring higher on either of these scales were more confident in their perceptual decisions. Trial-wise confidence judgements were expectedly driven by acoustic stimulus distances (i.e., the sensory evidence available), but also—to an equal magnitude—by LSHS scores thought to capture hallucination proneness (i.e., a perceptual prior or predisposition, see Fig. 6).

**Fig. 6.**
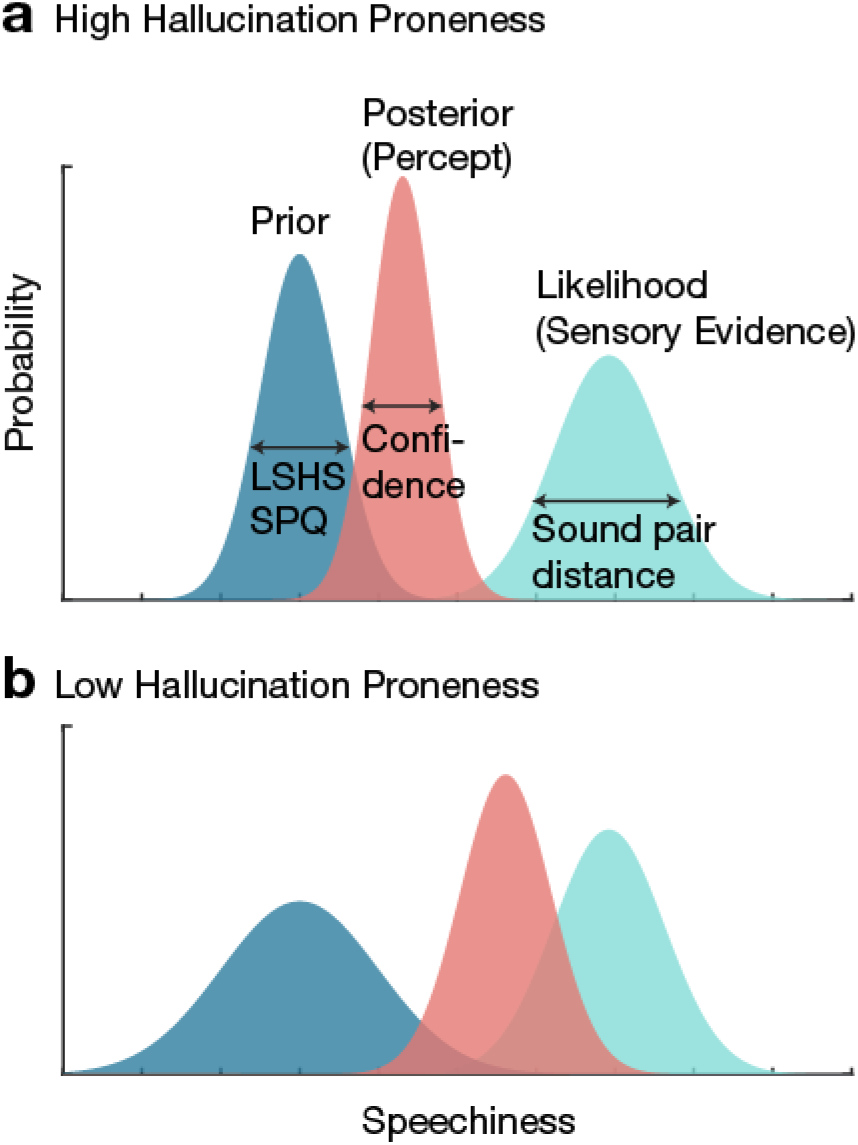
The current results in a framework of Bayesian models of perception. Perceptual inference in terms of the prior, likelihood (sensory evidence) and posterior (percept) is represented by Gaussian distributions over a perceptual dimension of ‘speechiness’; their widths represent precision. Both hallucination proneness and schizotypy (LSHS and SPQ) are thought to be proportional to the inverse width of the prior, while confidence being proportional to the inverse width of the posterior. Sensory evidence was operationalized as Euclidian sound distance, (a) In high hallucination-prone participants, the prior precision for a percept of ‘speechiness’ is thought to be higher (‘stronger prior’), contributing to the observed stronger confidence judgements (i.e., posterior precision) as compared to (b) low hallucination-prone individuals.

The present results are remarkably in line with an account of altered perceptual priors and a differential weighting of sensory evidence in hallucination-prone individuals, with accordingly changed perceptual decisions when classifying speech-like sounds.

### Speech perception in Hallucination proneness

The PCA analyses allowed us to examine to which degree different acoustic features were used for perceptual decision-making in varying levels of hallucination proneness. We found the presence of high-frequency components and – maybe more importantly – the absence of the speech-typical low temporal modulations to contribute to the classification into speech in hallucination-prone individuals. These effects were not specific to hallucination proneness but also observed in predisposition to delusions, emphasizing perceptual changes that seem to be shared in predisposition to both psychosis-like symptoms.

Hallucination proneness has been associated with more false alarms in auditory signal detection tasks. An increased false alarm rate was observed in a tone detection task with a conditioning visual stimulus in voice-hearers with and without psychosis^10^. In a speech in noise detection study, participants were to indicate “whether they had heard a voice”: Non-clinical hallucination-prone adults had more false alarms and expressed a liberal response bias ^40^. Yet, this study could not disentangle whether distinct acoustic features contribute to this bias towards a speech percept.

Why do the speechiness kernels in hallucination-prone individuals exhibit a high frequency dominance while being flat in the other two dimensions (i.e., the temporal and spectral modulations)? Our findings of atypical speech perception parallel the recent observations of differences in speech production in schizophrenia: aberrant acoustic patterns of vocal expressions (e.g., in pitch variability) have been reported alongside schizophrenia ^41^. The underlying cause common to aberrant speech perception and production may lie in a deficient auditory object formation as postulated for schizophrenia^23^, consistent with the notion that object formation relies on the intact extraction of spectro-temporal cues^24^. If auditory object formation is impaired, leading to typical speech frequencies being down-weighted, behaviourally a disadvantage would be expected for hallucination-prone individuals in challenging listening situations, e.g., when listening to competing speakers (at the ‘cocktail party’ ^42^).

Notably, the sound discriminability analysis showed no effects of hallucination proneness. This result indicates that incoming sensory evidence (operationalized as sound pair distance) was similarly amplified by speechiness kernels in hallucination-prone and control participants.

### Confidence Judgements in Hallucination proneness

In a predictive processing framework, the current results pose new evidence for changes to prior expectations in hallucination proneness ^14,43^: The data support an account of an increased precision or decreased variance in individual perceptual priors ^6,44^. The statistical model of single-trial confidence judgements in their own choices (“which [sound] is the more speech-like one?”) here provides important evidence.

First, as expected, the sensory evidence available on a given trial (i.e., width of an internal likelihood representation) exerts an impact on confidence. The expressed confidence is here thought of as the width or inverse precision of the posterior, reflecting the “noise” or uncertainty in one’s perceptual judgement^44^ (but see ^45^ for a different conceptualization). Accordingly, individual trait-like predispositions to perceive hallucinations can be considered stronger (i.e., less noisy and variable) perceptual priors, and they should hence contribute to a stronger confidence (i.e., smaller width of the posterior) in one’s own perceptual judgements, given that the incoming sensory evidence (i.e., likelihood) is equal. While this has been a guiding conjecture in the field of computational psychiatry (e.g.,^7^), it is borne out by the present evidence that hallucination proneness predicts confidence judgements. Figure 6 provides a schematic illustration of the evidence provided within a Bayesian model of perception. In sum, overly precise or strong perceptual priors that have previously been claimed for psychosis and hallucinations^5^ may contribute to an aberrant percept of ‘speechiness’.

The present results add novel evidence to a strain of findings supporting that prior beliefs mediate hallucinations ^5^. In a visual degraded image recognition task, individuals at risk of psychosis favored top-down prior knowledge of an image over available sensory evidence ^8^. Similarly, in the visual-auditory conditioning study by Powers, et al.^10^, voice-hearers experienced conditioned auditory hallucinations more often than control participants when seeing only the visual conditioning stimulus. Modelled in a Hierarchical Gaussian Filter^46^, this expressed as a higher weighting of prior beliefs over sensory evidence in voice-hearers ^10^. Note the different types of conceptualizations of priors, though: In the current study, explicit self-reports on the hallucination scales were thought of as a stable trait related to the width of the prior (Fig. 6). In contrast, Powers et al. derived the priors as model parameters as implicit estimate on a shorter task time scale.

An important open question remains at which neuronal levels these aberrant percepts emerge^14^. In contrast to tinnitus where usually simple tones or noises are perceived, AVH relate to the perception of complex sounds, that is, voices, in the absence of an external source. Similar to AVH ^5^, tinnitus has been proposed to be rooted in overly precise sensory evidence ^47^. In tinnitus, layer-specific effects in A1 have been postulated to lead to sharpening of a weak prior^47^.

In the genesis of AVH, computational changes in higher-level association cortex are likely^17^. For example, non-clinical voice hearers show enhanced fMRI responses in an executive attention network including cingulo-opercular and frontal cortex when listening to degraded (“sine-wave”) speech ^11^. Cingulo-opercular regions alongside with angular gyrus have been implicated in the semantic predictability gain ^48-50^ when rich semantic context informs an accurate prediction of upcoming speech under challenging listening conditions. In hallucination proneness, behavioural observations of increased semantic expectations^12^ suggest a bias towards such top-down information flow.

Top-down modulations from frontal to auditory cortex are known to be crucial for predicting the envelope during speech perception, where frontal signals causally influence the phase of brain oscillations in auditory cortex ^51^. Dysfunctional neural oscillations – which are thought to orchestrate neural responses throughout the brain – have been identified as candidate mechanism for a widespread network impairment in schizophrenia^52^.

However, it is also possible that in hallucination-prone individuals already the general response properties of auditory cortex are altered in terms of resting state activity (for review see ^14^), ongoing neural dynamics ^53^, spectral signatures ^54^, or encoding of spectro-temporal modulations^55^. The current study opens a specific and promising avenue using validated auditory perceptual-filtering models ^19,20,25^; Spectro-temporal modulations currently form a core tenet of auditory neuroscience, tractable for non-human animal research (e.g.,^56^) as well as human functional neuroimaging (e.g., ^22,55,57^)

In the present study, we tested populations where the overall level of hallucination proneness was nevertheless very modest (see Fig. 2). The speechiness kernel could easily be assessed in non-clinical and clinical frequent voice-hearers who may exhibit qualitatively similar but more extreme biases. This simple psychophysical paradigm presents a potential clinical instrument to improve prediction of conversion to psychosis in high-risk populations^4^.

### Conclusions

In sum, the current results endorse a continuum hypothesis of psychosis^3^ showing that individuals with different degrees of schizotypy – who sometimes experience auditory hallucinations but are not diagnosed with any psychotic disorder – do have distinct signatures of speech perception ^23^. Our results are remarkably in line with a Bayesian model of perception where stronger priors engender a bias towards hallucinations and foster perceptual confidence in light of ambiguous sensory input.

## Materials and methods

### Participants

#### Lab experiment

The final sample of the lab experiment comprised *N = 37* participants of which eleven were female; age ranged from 18 to 30 [mean 21.97] years. Exclusion criteria were self-reported hearing loss; neurological or psychiatric disorders; or the regular consumption of drugs, particularly amphetamines, cannabis, or similar psychoactive substances. Originally, *N* = 42 volunteers had been recruited, but data from five of these had to be excluded due to software problems during testing.

#### Online study

The final sample of the online study comprised *N* = 131 participants, of which 94 participants were female; age ranged from 19-61 [mean 27.2] years. A total of *N* = 353 volunteers had been recruited through dissemination of the study link in social media, university e-mail lists, and psychology student councils of different German Universities. *N* = 131 complete datasets were obtained. Self-exclusion was based on the same criteria as in the lab experiment.

All participants gave informed consent. Psychology students at the University of Lübeck were offered to obtain course credit. All procedures were approved by the local ethics committee of the University of Lübeck.

### Experimental Procedures

In the lab, participants first performed the psychoacoustic experiment, and then completed two questionnaires, namely the Schizotypal Personality Questionnaire (SPQ) and the Launay-Slade hallucination scale (LSHS). The rationale was to perform the relatively long psychoacoustic experiment first in order to not unnecessarily tire participants with the questionnaires before. Psychoacoustic testing was performed on a PC. Stimuli were delivered through Sennheiser HD 280 Pro headphones. Presentation level was kept constant at a comfortable and clearly audible level. In total, the duration of the lab experiment was approximately 60–75 minutes.

In the online experiment, the order of questionnaires and psychoacoustics was reversed, that is, participants first filled out the two questionnaires and afterwards performed the psychoacoustic experiment. The drop-out rate is expected to be higher in online than in lab experiments. Thus, the rationale for reversing the order was that participants dropping out after the questionnaires would at least provide SPQ and LSHS data. Also, the duration of the psychoacoustic online experiment was reduced to 20 % of duration of the lab experiment (speechiness kernels were shown to be highly similar with 100 and 540 trials, see supplementary Fig. S1), leaving the psychoacoustic testing shorter and less tiring. The order of the two questionnaires was randomized across participants.

For the psychoacoustic task, participants were instructed to use headphones. Prior to the psychoacoustic task, participants could adjust the presentation intensity to a comfortable level using an exemplar sound texture and were instructed to keep the presentation level constant during the experiment. The total duration of the experiment amounted to approximately 20 - 25 min. Participants were debriefed after the experiment.

### Questionnaires

Schizotypy was assessed using the German adaptation of the schizotypal personality questionnaire (SPQ ^29,30^). The SPQ comprises nine subscales based on *DSM-III-R* criteria for diagnosis of schizotypal personality disorder, namely: ideas of reference; excessive social anxiety; odd beliefs or magical thinking; unusual perceptual experiences; odd or eccentric behavior; no close friends; odd speech; constricted affect; and suspiciousness. In the current study, we were particularly interested in the subscale ‘unusual perceptual experiences’ (SPQ-UP) to measure predisposition to hallucinations. In total, the SPQ includes 74 items of which the responses (true/false) are summed up to derive a total score, amounting to a maximal total score of 74. The SPQ-UP subscale has a maximum total score of nine.

Predisposition for hallucinations was assessed using the German version of the Launay-Slade Hallucination Scale (LSHS ^31,32^). The questionnaire comprises twelve items which are assessed on a five-point Likert scale (0 – 4). The LSHS score is derived as the sum of all items and can thus maximally reach 48.

### Psychoacoustic testing

#### Stimuli

Stimuli were resynthesized natural sounds (“sound textures”^26^) presented in white noise at an SNR of 3 dB. Textures were synthesized from the spectro-temporal modulation content of a large set of real-life sounds (*n* = 192), including speech, voice, animal vocalizations as well as nature and tool (instrument) sounds we had used in a previous study^21^ (see Fig. 1a,b). Texture synthesis parameters were as follows: frequency range 0.02–10 kHz, number of frequency bands = 30, sampling rate = 20 kHz, temporal modulation range 0.5–200 Hz, sampling rate = 400 Hz; maximum number of iterations = 60. Textures had a length of 1 s and a sampling rate of 20 kHz. Out of the total of 192 textures, we selected the 108 “most speech-like” textures for the final experiment: “speech-like” textures were defined as those textures whose spectral centroid diverted less than 3 standard deviations from the mean spectral centroid of those textures that had originally been synthesized from speech stimuli.

#### Task

Two sounds were randomly paired on each trial and presented with an inter-stimulus-interval of 500 ms. In a two-alternative-forced-choice (2-AFC) task, participants were asked to decide “which exemplar sounded more like speech” (Fig. 1c). Participants rated their confidence from 1-3 (unconfident to confident).

The lab experiment was performed in Matlab 2018b. The order of exemplars was randomized across participants. In total, each texture exemplar was presented five times. The first ten trials were training trials. Twenty relatively unambiguous catch trials were distributed evenly across the experiment. In both catch and training trials, a speech texture (that is, a texture of which the spectral centroid diverted < 0.6 from the mean spectral centroid of speech textures) was paired with a texture from the other categories. In total, the lab experiment comprised 560 trials, including 20 catch trials. Catch and training trials were excluded from the subsequent reverse correlation analyses, such that speechiness kernels were estimated based on 540 trials.

The online experiment was identical to the lab experiment with the following exceptions: the online experiment was performed in Labvanced (Scicovery GmbH, Osnabrück, Germany). The order of exemplars was fixed across participants due to programming constraints in Labvanced. To keep the experiment short (approximately 10 min), each texture exemplar was presented only once. The first four trials were training trials. Five relatively unambiguous catch trials were distributed evenly across the experiment. In total, the online experiment comprised 117 trials out of which of 5 were catch trials. Catch and training trials were excluded from the subsequent reverse correlation analyses, such that speechiness kernels were estimated based on 108 trials. Note that kernels estimated based on 100 trials are highly similar to the ones based on 540 trials, but for details on kernel stability see supplementary Fig. S1.

### Analyses

#### Sound decomposition

We analyzed how participants’ judgements varied as a function of the spectro-temporal modulation content of the stimuli. The modulation content of the stimuli was obtained by filtering the sounds with a model of auditory processing ^20^ using the “NSL Tools” package (available at http://www.isr.umd.edu/Labs/NSL/Software.htm) and customized Matlab code (The MathWorks Inc., Matlab 2014b/2018a). First, spectrograms for all sounds were obtained using a bank of 128 overlapping bandpass filters with equal width (Q_10dB_ = 3), spaced along a logarithmic frequency axis over a range of *f* = 116–2872 Hz (hair cell stage). A midbrain stage modelled the enhancement of frequency selectivity as a first-order derivative with respect to the frequency axis, followed by a half-wave rectification and a short-term temporal integration (time constant *τ* = 8 ms).

Then, the auditory spectrogram was analyzed by the cortical stage, where the modulation content of the auditory spectrogram was computed through a bank of 2-dimensional filters selective for a combination of spectral and temporal modulations. The filter bank performs a complex wavelet decomposition of the auditory spectrogram. The magnitude of such decomposition yields a phase-invariant measure of modulation content. The modulation selective filters have joint selectivity for spectral and temporal modulations, and are directional, i.e. they respond either to upward or downward frequency sweeps.

#### Modulation filters

Filters were tuned to spectral modulation frequencies of Ω = [0.5, 0.76, 1.15, 1.74, 2.64, 4] cyc/oct, temporal modulation frequencies of ω = [1, 2, 4, 8, 16, 32] Hz, and centre frequencies of *f* = [116, 183, 290, 459, 726, 1148, 1816, 2872] Hz. The filter bank output was computed at each frequency along the tonotopic axis and then averaged over time. The time-averaged output of the filter bank was averaged across the upward and downward filter directions. This resulted in a representation with 6 spectral modulation frequencies, 6 temporal modulation frequencies, and 8 tonotopic frequencies, amounting to 288 acoustic features in total. The rationale for this choice of values was to use a decomposition roughly covering the temporal and spectral modulations present in speech (for spectro-temporal modulation content of all sound categories see Fig. 1a, for modulation content of speech see Fig. 1b).

#### Psychophysical reverse correlation

To obtain an internal template of speech, we used the reverse correlation technique ^58,59^. Sounds were sorted into speech and nonspeech stimuli based on the participants’ responses. Note that due to the 2-AFC task, the two choices are symmetric, that is, on each trial one stimulus was attributed to the speech and hence the other one to the nonspeech category. Therefore, to obtain individual speechiness kernels we subtracted the two templates for speech and nonspeech:

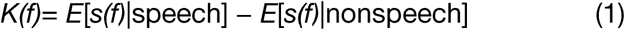

where *E*[*s*(*f*)|speech] indicates the trial average of the stimulus at feature *f* conditional on choice “speech”, *s(f)* is the stimulus at feature *f*, and *K(f)* is the magnitude of the psychophysical kernel at feature *f*.

#### Kernel stability

To obtain an estimate of how many trials are necessary to obtain a stable estimate of individual speechiness kernels, we assessed kernel stability in the lab-experiment data. To this end, we iteratively calculated the Pearson’s correlation between the kernel based on a subset of *n* trials and the final kernel based on all 540 trials^60^ for all participants.

#### Principal component analysis (PCA)

To reduce dimensionality of the speechiness kernel (comprising *n* = 288 acoustic features), we computed the matrix singular value decomposition on the set of all lab and online speechiness kernels (all scaled between 0 and 1) jointly in Matlab 2018a and jamovi 1.0.7.0. This yielded *k* = 288 components. We decided on the number of components to retain based on a scree plot ^61^ and parallel analysis ^62^. We retained only those components where the eigenvalues associated with the raw data exceed the eigenvalues of surrogate data, leaving us with the first six components (see scree plot Fig. 4b). The individual speechiness kernels were then projected through (i.e., multiplied with) those components. The individual component loadings were scaled by the singuiar values, yielding individual component scores. Lastly, the component scores were correlated using Pearson’s *r* with the LSHS and SPQ-UP subscale, respectively. The two scales (LSHS and SPQ-UP subscale) were chosen as they measure psychotic-like experiences, specifically predisposition for unusual perceptual experiences (e.g., hallucinations).

#### Sound discriminability analysis

To estimate the discriminability of two textures presented on a trial, we first calculated trial-wise sound pair distance as the Euclidian distance between each of the 288 features. To investigate whether individual speechiness kernels influenced the discriminability of sound pairs presented on each trial, sounds in the modulation space (comprising 288 features) were projected through (multiplied with) individual speechiness kernels. Then, for each sound separately, the features were normalized between their minimum and maximum (effectively scaling them between 0 and 1). Sound pair distance was compared before and after projection through speechiness kernels.

#### Generalized linear mixed-effects models

To evaluate the relation of trial-wise confidence judgements with sound-pair discriminability (see above) and hallucination proneness, we used a Cumulative Link Mixed Models, that is, a hierarchical generalized linear model with a cumulative link function (i.e., an ordinal regression model). Ordinal linear mixed-effects regression was fitted using the Bayesian estimation package *brms* in RStudio 1.3.959 (cumulative-probit link function). We regressed trial-by-trial confidence judgements (on a Likert scale from 1 [“not confident”] to 3 [“confident”]) against the z-scored predictors trial-wise sound pair distance, LSHS score, global SPQ score, as well as experiment type (online, lab) as covariate of no interest with subject-specific random intercepts.

## Acknowledgements

This research was funded by an ERC consolidator Grant (ERC-CoG-2014-646696 “AUDADAPT” to J.O.). We thank Felix Greuling for the implementation of the online study and Tabea Landschoff, Franziska Mauz, Hannah Kleinhaus for data acquisition.

## References

1 de Leede-Smith, S. & Barkus, E. A comprehensive review of auditory verbal hallucinations: lifetime prevalence, correlates and mechanisms in healthy and clinical individuals. Frontiers in human neuroscience 7, 367, doi: 10.3389/fnhum.2013.00367 (2013).

2 Linscott, R. J. & van Os, J. An updated and conservative systematic review and meta-analysis of epidemiological evidence on psychotic experiences in children and adults: on the pathway from proneness to persistence to dimensional expression across mental disorders. Psychological medicine 43, 1133–1149, doi: 10.1017/S0033291712001626 (2013).

3 Sommer, I. E. et al. Healthy individuals with auditory verbal hallucinations; who are they? Psychiatric assessments of a selected sample of 103 subjects. Schizophrenia bulletin 36, 633–641, doi:10.1093/schbul/sbn130 (2010).

4 Lehembre-Shiah, E. et al. Distinct Relationships Between Visual and Auditory Perceptual Abnormalities and Conversion to Psychosis in a Clinical High-Risk Population. JAMA psychiatry 74, 104–106, doi: 10.1001/jamapsychiatry.2016.3055 (2017).

5 Corlett, P. R. et al. Hallucinations and Strong Priors. Trends Cogn Sci 23, 114–127, doi:10.1016/j.tics.2018.12.001 (2019)

6 Sterzer, P. et al. The Predictive Coding Account of Psychosis. Biological psychiatry 84, 634–643, doi:10.1016/j.biopsych.2018.05.015 (2018).

7 Adams, R. A., Huys, Q. J. & Roiser, J. P. Computational Psychiatry: towards a mathematically informed understanding of mental illness. Journal of neurology, neurosurgery, and psychiatry 87, 53–63, doi:10.1136/jnnp-2015-310737 (2016).

8 Teufel, C. et al. Shift toward prior knowledge confers a perceptual advantage in early psychosis and psychosis-prone healthy individuals. Proceedings of the National Academy of Sciences of the United States of America 112, 13401–13406, doi:10.1073/pnas.1503916112 (2015).

9 Davies, D. J., Teufel, C. & Fletcher, P. C. Anomalous Perceptions and Beliefs Are Associated With Shifts Toward Different Types of Prior Knowledge in Perceptual Inference. Schizophrenia bulletin 44, 1245–1253, doi:10.1093/schbul/sbx177 (2018).

10 Powers, A. R., Mathys, C. & Corlett, P. R. Pavlovian conditioning-induced hallucinations result from overweighting of perceptual priors. Science 357, 596–600, doi:10.1126/science.aan3458 (2017).

11 Alderson-Day, B. et al. Distinct processing of ambiguous speech in people with non-clinical auditory verbal hallucinations. Brain: a journal of neurology 140, 2475–2489, doi:10.1093/brain/awx206 (2017).

12 Vercammen, A. & Aleman, A. Semantic expectations can induce false perceptions in hallucination-prone individuals. Schizophrenia bulletin 36, 151156, doi:10.1093/schbul/sbn063 (2010).

13 Daalman, K., Verkooijen, S., Derks, E. M., Aleman, A. & Sommer, I. E. The influence of semantic top-down processing in auditory verbal hallucinations. Schizophrenia research 139, 82–86, doi:10.1016/j.schres.2012.06.005 (2012).

14 Horga, G. & Abi-Dargham, A. An integrative framework for perceptual disturbances in psychosis. Nature reviews. Neuroscience 20, 763–778, doi: 10.1038/s41583-019-0234-1 (2019).

15 Horga, G., Schatz, K. C., Abi-Dargham, A. & Peterson, B. S. Deficits in predictive coding underlie hallucinations in schizophrenia. The Journal of neuroscience: the official journal of the Society for Neuroscience 34, 8072–8082, doi:10.1523/JNEUROSCI.0200-14.2014 (2014).

16 Dierks, T. et al. Activation of Heschl’s gyrus during auditory hallucinations. Neuron 22, 615–621 (1999).

17 Allen, P., Laroi, F., McGuire, P. K. & Aleman, A. The hallucinating brain: a review of structural and functional neuroimaging studies of hallucinations. Neuroscience and biobehavioral reviews 32, 175–191, doi: 10.1016/j.neubiorev.2007.07.012 (2008).

18 Formisano, E. et al. Mirror-symmetric tonotopic maps in human primary auditory cortex. Neuron 40, 859–869 (2003).

19 Dau, T., Kollmeier, B. & Kohlrausch, A. Modeling auditory processing of amplitude modulation. I. Detection and masking with narrow-band carriers. The Journal of the Acoustical Society of America 102, 2892–2905 (1997).

20 Chi, T., Ru, P. & Shamma, S. A. Multiresolution spectrotemporal analysis of complex sounds. The Journal of the Acoustical Society of America 118, 887–906 (2005).

21 Erb, J. et al. Homology and Specificity of Natural Sound-Encoding in Human and Monkey Auditory Cortex. Cerebral cortex 29, 3636–3650, doi:10.1093/cercor/bhy243 (2019).

22 Santoro, R. et al. Reconstructing the spectrotemporal modulations of real-life sounds from fMRI response patterns. Proceedings of the National Academy of Sciences of the United States of America 114, 4799–4804, doi: 10.1073/pnas.1617622114 (2017).

23 Uhlhaas, P. J. & Mishara, A. L. Perceptual anomalies in schizophrenia: integrating phenomenology and cognitive neuroscience. Schizophrenia bulletin 33, 142–156, doi:10.1093/schbul/sbl047 (2007).

24 Bizley, J. K. & Cohen, Y. E. The what, where and how of auditory-object perception. Nature reviews. Neuroscience 14, 693–707, doi:10.1038/nrn3565 (2013).

25 Elliott, T. M. & Theunissen, F. E. The modulation transfer function for speech intelligibility. PLoS computational biology 5, e1000302, doi:10.1371/joumal.pcbi.1000302 (2009).

26 McDermott, J. H. & Simoncelli, E. P. Sound texture perception via statistics of the auditory periphery: evidence from sound synthesis. Neuron 71, 926–940, doi:10.1016/j.neuron.2011.06.032 (2011).

27 Murray, R. F. Classification images: A review. Journal of vision 11, doi:10.1167/11.5.2 (2011).

28 Ahumada, A. & Lovell, J. Stimulus features in signal detection. The Journal of the Acoustical Society of America 49, 1751–1756 (1971).

29 Raine, A. et al. Cognitive-perceptual, interpersonal, and disorganized features of schizotypal personality. Schizophrenia bulletin 20, 191–201, doi: 10.1093/schbul/20.1.191 (1994).

30 Klein, C., Andresen, B. & Jahn, T. Erfassung der schizotypen Persönlichkeit nach DSM-III-R: Psychometrische Eigenschaften einer autorisierten deutschsprachigen Übersetzung des Schizotypal Personality Questionnaire” (SPQ) von Raine. Diagnostica 43, 347–369, doi:https://doi.org/10.1037/t10727-000 (1997).

31 Launay, G. & Slade, P. The measurement of hallucinatory predisposition in male and female prisoners.. Personality and Individual Differences 2, 221–234, doi:https://doi.org/10.1016/0191-8869(81)90027-1 (1981).

32 Lincoln, T. M., Keller, E. & Rief, W. Die Erfassung von Wahn und Halluzinationen in der Normalbevölkerung: Deutsche Adaptatíonen des Peters et al. Delusions Inventory (PDI) und der Launay Slade Hallucination Scale (LSHS-R). Diagnostica 55, 29–40, doi:https://doi.org/10.1026/0012-1924.55.1.29 (2009).

33 Laroì, F. How do auditory verbal hallucinations in patients differ from those in non-patients? Frontiers in human neuroscience 6, 25, doi: 10.3389/fnhum.2012.00025 (2012).

34 Giraud, A. L. & Pœppel, D. Cortical oscillations and speech processing: emerging computational principles and operations. Nature neuroscience 15, 511–517, doi:10.1038/nn.3063 (2012).

35 Ding, N., Melloni, L., Zhang, H., Tian, X. & Pœppel, D. Cortical tracking of hierarchical linguistic structures in connected speech. Nature neuroscience 19, 158–164, doi:10.1038/nn.4186 (2016).

36 Gross, J. et al. Speech rhythms and multiplexed oscillatory sensory coding in the human brain. PLoS Biol 11, e1001752, doi:10.1371/journal.pbio.1001752 (2013).

37 Holland, P. W. & Weisch, R. E. Robust regression using iteratively reweighted least-squares. Communications in Statistics - Theory and Methods 6, 813–827, doi:10.1080/03610927708827533 (1977).

38 Gaweda, L. & Moritz, S. The role of expectancies and emotional load in false auditory perceptions among patients with schizophrenia spectrum disorders. European archives of psychiatry and clinical neuroscience, doi: 10.1007/s00406-019-01065-2 (2019).

39 Hoven, M. et al. Abnormalities of confidence in psychiatry: an overview and future perspectives. Translational psychiatry 9, 268, doi:10.1038/s41398-019-0602-7 (2019).

40 Barkus, E. et al. Auditory false perceptions are mediated by psychosis risk factors. Cognitive neuropsychiatry 16, 289–302, doi:10.1080/13546805.2010.530472 (2011).

41 Parola, A., Simonsen, A., Bliksted, V. & Fusaroli, R. Voice patterns in schizophrenia: A systematic review and Bayesian meta-analysis. Schizophrenia research 216, 24–40, doi:10.1016/j.schres.2019.11.031 (2020).

42 Ding, N. & Simon, J. Z. Emergence of neural encoding of auditory objects while listening to competing speakers. Proceedings of the National Academy of Sciences of the United States of America 109, 11854–11859, doi: 10.1073/pnas.1205381109 (2012).

43 Teufel, C. & Fletcher, P. C. Forms of prediction in the nervous system. Nature reviews. Neuroscience 21, 231–242, doi:10.1038/s41583-020-0275-5 (2020).

44 Adams, R. A., Stephan, K. E., Brown, H. R., Frith, C. D. & Friston, K. J. The computational anatomy of psychosis. Frontiers in psychiatry 4, 47, doi: 10.3389/fpsyt.2013.00047 (2013).

45 Pouget, A,, Drugowitsch, J. & Kepecs, A. Confidence and certainty: distinct probabilistic quantities for different goals. Nature neuroscience 19, 366–374, doi:10.1038/nn.4240 (2016). “

46 Mathys, C. D. et al. Uncertainty in perception and the Hierarchical Gaussian Filter. Frontiers in human neuroscience 8, 825, doi:10.3389/fnhum.2014.00825 (2014).

47 Sedley, W., Friston, K. J., Gander, P. E., Kumar, S. & Griffiths, T. D. An Integrative Tinnitus Model Based on Sensory Precision. Trends Neurosci 39, 799–812, doi:10.1016/j.tins.2016.10.004 (2016).

48 Rysop, A. U., Schmitt, L. M., Obleser, J. & Hartwigsen, G. Neural modelling of the semantic predictability gain under challenging listening conditions. Hum ßra/n Mapp, doi:10.1002/hbm.25208 (2020).

49 Obleser, J., Wise, R. J., Dresner, M. A. & Scott, S. K. Functional integration across brain regions improves speech perception under adverse listening conditions. The Journal of neuroscience: the official journal of the Society for Neuroscience 27, 2283–2289, doi:10.1523/JNEUROSCI.4663-06.2007 (2007).

50 Obleser, J. & Kotz, S. A. Expectancy constraints in degraded speech modulate the language comprehension network. Cerebral cortex 20, 633–640, doi: 10.1093/cercor/bhp128 (2010).

51 Park, H., Ince, R. A., Schyns, P. G., Thut, G. & Gross, J. Frontal top-down signals increase coupling of auditory low-frequency oscillations to continuous speech in human listeners. Current biology: CB 25, 1649–1653, doi: 10.1016/j.cub.2015.04.049 (2015).

52 Uhlhaas, P. J. & Singer, W. Abnormal neural oscillations and synchrony in schizophrenia. Nature Reviews Neuroscience 11, 100–113, doi:10.1038/nrn2774 (2010).

53 Waschke, L, Tune, S. & Obleser, J. Local cortical desynchronization and pupil - linked arousal differentially shape brain states for optimal sensory performance. eLife 3, doi:10.7554/eLife.51501 (2019).

54 Keitel, A. & Gross, J. Individual Human Brain Areas Can Be Identified from Their Characteristic Spectral Activation Fingerprints. PLoS Biol 14, e1002498, doi:10.1371/journal.pbio.1002498 (2016).

55 Erb, J., Schmitt, L. M. & Obleser, J. Temporal selectivity declines in the aging human auditory cortex. eLife 9, doi:10.7554/eLife.55300 (2020).

56 Baumann, S. et al. The topography of frequency and time representation in primate auditory cortices. eLife 4, doi:10.7554/eLite.03256 (2015).

57 Santoro, R. et al. Encoding of natural sounds at multiple spectral and temporal resolutions in the human auditory cortex. PLoS computational biology 10, e1003412, doi:10.1371/journal.pcbi.1003412 (2014).

58 Neri, P. & Heeger, D. J. Spatiotemporal mechanisms for detecting and identifying image features in human vision. Nature neuroscience 5, 812–816, doi:10.1038/nn886 (2002).

59 de Beer, R. & Kuyper, P. Triggered correlation. IEEE transactions on bio-medical engineering 15, 169–179, doi:10.1109/tbme.1968.4502561 (1968).

60 Burred, J. J., Ponsot, E., Goupil, L, Liuni, M. & Aucouturier, J. J. CLEESE: An open-source audio-transformation toolbox for data-driven experiments in speech and music cognition. PloS one 14, e0205943, doi:10.1371/journal.pone.0205943 (2019).

61 Cattell, R. B. The Scree Test For The Number Of Factors. Multivariate behavioral research 1, 245–276, doi:10.1207/s15327906mbr0102_10 (1966).

62 Horn, J. L. A Rationale and Test for the Number of Factors in Factor Analysis. Psychometrika 30, 179–185, doi:10.1007/BF02289447 (1965).

