## Supplementary Figures for "Aberrant perceptual judgements on speech-relevant acoustic features in hallucination-prone individuals"

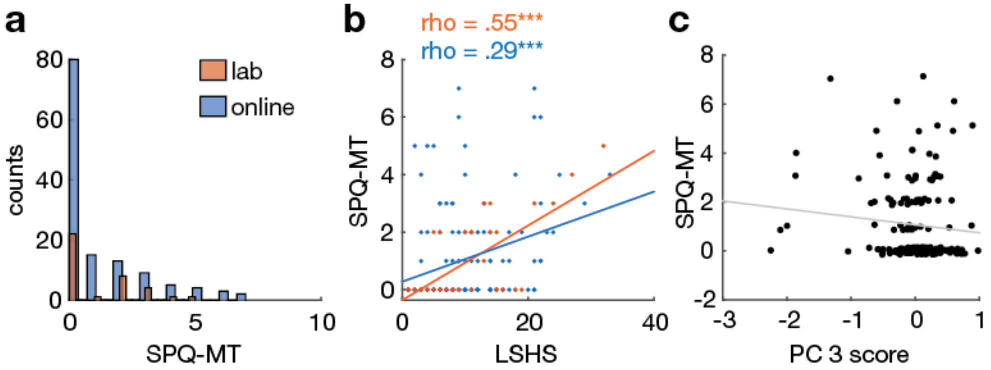


**Figure S1. SPQ subscale Magical Thinking and its relation to LSHS and the Speechiness kernel.** (a) Histograms for SPQ-MT scores separately for lab (orange) and online (blue) experiment. Note that the maximum possible subscale score is 7 for SPQ-MT. (b) Scatter plot showing the correlation between SPQ-UP and LSHS scores separately for lab (red) and online experiment (blue). (c) Scatter plot showing the correlation between SPQ-UP scores and the scores of the third principal component of the Speechiness kernel. Pearson's correlation cœfficient $r$ = -0.103, $p$ = 0.185, Spearman's correlation cœfficient $rho$ = -0.142, $p$ = 0.067, Mutual Information $MI$ = -0.006, $p$ = 0.125. For display purposes only the integer SPQ-UP scores were jittered slightly. LSHS: Launay-Slade hallucination scale; SPQ-MT: schizotypal personality questionnaire subscale Magical Thinking. *** $p < .001$.

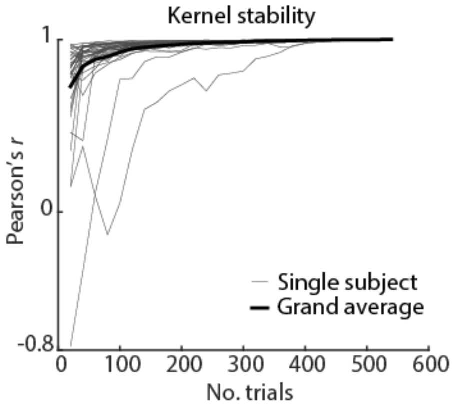


**Figure S2. Stability of speechiness kernels in the lab experiment.** Correlation of speechiness kernels with the final kernel (based on all trials) as a function of the numbers of trials used to estimate the kernel. On average, speechiness kernels based on the first 100 trials were highly similar to the final kernel, which we took as evidence that 108 trials are sufficient for the online experiment to obtain a reliable estimate of the speechiness kernel.

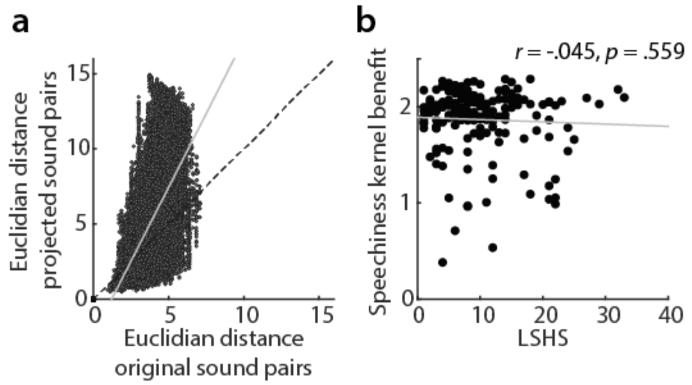


**Figure S3. Effect of speechiness kernels on discriminability of sound pairs.** (a) We filtered all sounds (presented across experiments and participants, n = 34,128 sound pairs) by individual speechiness kernels, leading overall to higher discriminability (euclidian distance in the modulation space) of projected compared to original sound pairs. (b) In this space, we fitted individual linear regression lines. The slope of this linear fit ('speechiness kernel benefit') was above 1 for most participants, indicating that filtering with individual speechiness kernels improved discriminability of sound pairs. However, this benefit was unrelated to hallucination proneness, as shown by Pearson's r. LSHS: Launay-Slade hallucination scale.
